# Cdc42 Partitioning by Chaperone Ydj1 During Asymmetric Division and Aging in Yeast

**DOI:** 10.1101/2025.07.10.664052

**Authors:** Pil Jung Kang, Hana Mazak, Sung Sik Lee, Hay-Oak Park

## Abstract

Cdc42, a small GTPase crucial for cell polarity, often becomes hyperactive with age and contributes to senescence and aging in yeast and animal cells. However, the mechanisms underlying its age-related upregulation are not well understood. Here, we report that in budding yeast, Cdc42 accumulates over successive divisions and that lowering its levels can extend lifespan. Using microfluidics-assisted live-cell imaging and genetic analysis, we found that Cdc42 is distributed unevenly between mother and daughter cells during division. Daughter cells inherit lower levels of Cdc42, which likely helps them remain young. This asymmetric distribution depends on Cdc42’s localization to endomembranes and involves Ydj1, a farnesylated Hsp40/DnaJ chaperone anchored to the <u>endoplasmic reticulum (**ER**</u>). Ydj1 interacts with Cdc42, enhancing its stability and proper partitioning during cell division. We thus propose that ER-bound Ydj1 facilitates the asymmetric distribution of Cdc42, limiting aging to mother cells. We thus propose that ER-bound Ydj1 facilitates the asymmetric distribution of Cdc42, limiting aging to mother cells.

## INTRODUCTION

Cdc42 is crucial for establishing cell polarity by regulating cytoskeletal organization and directional growth in organisms ranging from yeast to humans [1, 2]. Although it is essential for normal cell function, increased Cdc42 activity in aged cells—such as yeast and hematopoietic stem cells—results in polarity loss and more symmetric division, but the mechanisms behind this are not fully understood [3–7]. In budding yeast, polarity is essential for asymmetric cell division, which leads to mother cell aging, while daughter cells retain their full lifespan potential. The number of divisions a mother cell undergoes before becoming senescent defines its <u>replicative lifespan (hereafter, **lifespan**)</u> [8]. While the number of senescent cells increases with age in multicellular organisms [9], senescence is more directly linked to limited lifespan in yeast. Aging-associated cellular features, such as increased protein aggregates [10–12], mitochondrial dysfunction [13, 14], and loss of rDNA silencing, leading to <u>extrachromosomal</u> <u>rDNA circles (**ERCs**)</u> [15, 16], are generally restricted to mother cells in yeast and also relate to metazoan aging [17, 18]. Sir2 silences the rDNA locus and suppresses ERC formation in yeast [19], whereas Fob1 destabilizes rDNA and promotes ERC production [20]. Therefore, deletion of *SIR2* shortens lifespan, while deletion of *FOB1* extends it compared to <u>wild type (**WT**)</u> [19, 21].

During polarized growth in budding yeast, the ‘polarisome’—a multi-protein complex crucial for establishing and maintaining cell polarity [22]—facilitates retrograde transport of protein aggregates from buds to mother cells along actin cables [23]. This process also requires Sir2, which promotes proper folding of actin through the chaperonin CCT [23]. Yet, mutations in polarisome components do not completely abolish asymmetric segregation of protein aggregates [24, 25], indicating that additional mechanisms contribute to mother-cell-specific aging.

Yeast cells lacking Cdc42-inhibitory polarity cues or Rga1, a Cdc42 GTPase-activating protein (GAP), display shortened lifespans [7, 26]. Our previous microfluidic imaging with a Cdc42-GTP biosensor revealed hyperactivation of Cdc42 during aging. Mild overexpression of Cdc42, while not hindering exponential growth, shortens lifespan and exhibits characteristics of premature aging [7]. Notably, *rga1Δ* cells, which have elevated Cdc42-GTP levels even at young ages, show further increase in Cdc42-GTP as they age. Similarly, <u>cells overexpressing *CDC42* (*CDC42*</u>*<u>_OV_</u>*) display a wide range of Cdc42 increases with aging [7]. These observations led us to examine how Cdc42 protein levels are regulated during aging.

Here, we report that Cdc42 accumulates with age and is unevenly segregated between mother and daughter cells during division. This asymmetric distribution, as well as maintenance of Cdc42 levels, depends on its endomembrane association and interaction with Ydj1, a farnesylated Hsp40/DnaJ chaperone. Our findings highlight a critical role for ER-bound Ydj1 in the unequal partitioning of Cdc42 during asymmetric division and likely during aging.

## RESULTS AND DISCUSSION

### Cdc42 levels increase in aged cells, likely limiting lifespan

Given that cells lacking the Cdc42 GAP Rga1 show increased Cdc42 activation as they age [7], we investigated whether Cdc42 protein levels change across successive cell divisions. We also sought to understand how daughter cells maintain low Cdc42 levels despite strong polarization to the bud tip early in the cell cycle [27]. Using time-lapse imaging of cells expressing Cdc42-mCherry^SW^ (a fully functional internal fusion [28]) and Cdc3-GFP (a septin marking mother and bud compartments), we observed that Cdc42-mCherry^SW^ fluorescence peaked in small buds early in the cell cycle and declined as the bud grew, reaching a minimum in large buds or newborn daughter cells (S1_Fig). We therefore focused subsequent analyses on cells in late M phase or cytokinesis, when Cdc42 no longer polarizes to the bud and before it polarizes to a new bud site.

To compare Cdc42 levels at different ages, we performed microfluidic imaging of WT cells expressing Cdc42-mCherry^SW^ (Fig. 1A; S1_Movie) and measured mean fluorescence intensity in each cell during its first three divisions (‘young mother’) and last three divisions before death (‘old mother’). These analyses showed that Cdc42 levels increase with cell age (Fig. 1B). We next examined how reduced Cdc42 protein levels affect lifespan using the *cdc42-108(R147A E148A K150A)* allele. This hypomorphic variant produces approximately 50% less Cdc42 protein than the isogenic WT (Fig. 1C) but is phenotypically indistinguishable from WT during exponential growth [29]. Lifespan analysis using microfluidic imaging showed that *cdc42-108* cells had a median lifespan about 23.5% longer than WT (Fig. 1D). To determine whether this extended lifespan was due to reduced Cdc42 protein rather than loss of function, we introduced an extra copy of *cdc42-108* into the mutant. This strain, carrying two copies of the *cdc42-108* allele, had a lifespan similar to WT and shorter than the single-copy *cdc42-108* mutant (Fig. 1E). Taken together with the shorter lifespan of the *CDC42_OV_* strain [7], these results suggest that increased Cdc42 levels likely limit lifespan.

**Fig. 1.**
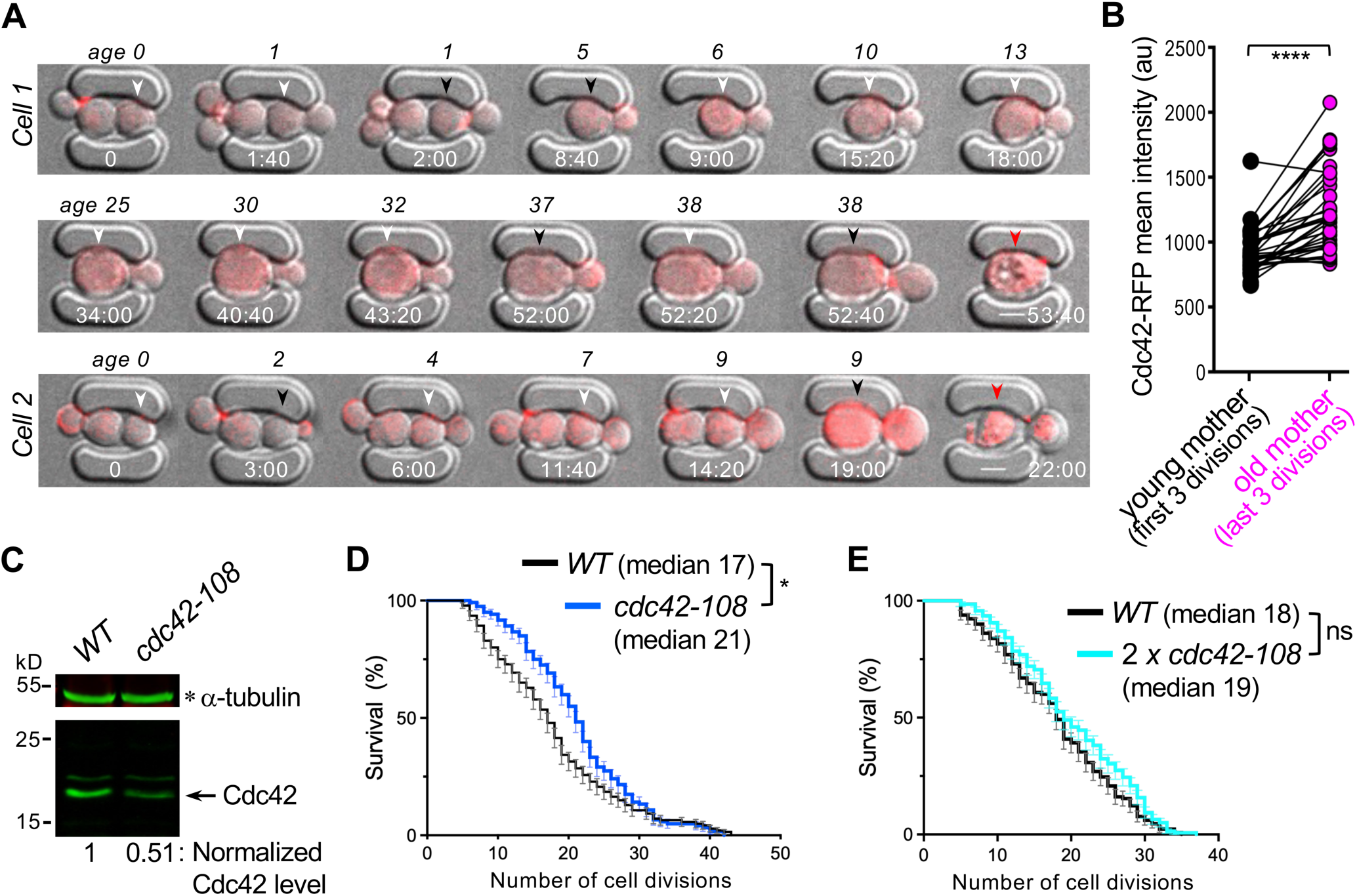
Cdc42 accumulates in aged cells, and reducing its levels extends lifespan **A.** Two representative WT cells expressing Cdc42-mCherry^SW^ are shown at selected ages and time points (hr: min) from initial loading. White arrowheads indicate cells during or shortly after cytokinesis, when Cdc42 levels are quantified. Black and red arrowheads denote the same cells at other cell cycle stages and cell death, respectively. Scale bar: 3 µm. See S1_Movie. **B.** Mean fluorescence intensity of Cdc42-mCherry^SW^ in mother cells at young (first three divisions) vs. old (last three divisions) ages, measured at the large-budded stage (see Fig. 1A legend). Each circle represents the average intensity from three divisions of an individual cell. n = 37; ****, *p* < 0.0001, paired t-tests. **C.** Cdc42 proteins in WT and *cdc42-108* cells, grown at 30°C to mid-log phase, were detected by immunoblotting using a monoclonal anti-Cdc42 antibody, and their levels were normalized to the loading control, α-tubulin. **D.** Percentage of cell survival (mean ± SEM) at each age of WT and *cdc42-108* mutant. Median lifespan is shown. n = 140 (WT) and 120 (*cdc42-108*); *, *p* = 0.0289, Log-Rank test. **E.** Cell survival percentage (mean ± SEM) of WT and a strain with two copies of *cdc42-108* (2 x *cdc42-108*). Median lifespan is indicated. n = 130 (WT) and 139 (*2 x cdc42-108*); ns (not significant), *p* = 0.063, Log-Rank test.

### Upregulation of Cdc42 likely limits lifespan independently of rDNA silencing loss

Cdc42 is mainly known for its role in establishing polarity at the cell cortex, but it also localizes to intracellular sites associated with the nuclear envelope and vacuoles [28, 30]. Since ERCs associate with the nuclear pore complex and accumulate asymmetrically in mother cells [31, 32], we examined whether increased Cdc42 could promote aging by interfering with rDNA silencing. Previous studies have shown that distinct terminal bud shapes are linked to specific aging defects: elongated buds correlate with rDNA desilencing and nucleolar destabilization [33, 34], while round buds are associated with mitochondrial dysfunction [35, 36]. In WT cells near death, elongated buds appear in approximately 20% of cases, rising to about 60% in *sir2Δ* mutants [37, 38], supporting a connection between silencing defects and elongated bud morphology.

Our microfluidic imaging of WT cells in two genetic backgrounds revealed that most aging cells displayed round buds or no buds at terminal stages (Fig. 2A). Similarly, elongated buds were uncommon in *CDC42ov* cells, suggesting that elevated Cdc42 does not promote loss of rDNA silencing. To further test whether Cdc42 upregulation influences rDNA silencing, we examined the effects of the *cdc42-108* mutation in a *fob1Δ* strain, a long-lived mutant with reduced ERC formation [21]. Remarkably, the *cdc42-108 fob1Δ* double mutant showed about a 21% increase in median lifespan compared to *fob1Δ* alone (Fig. 2B), indicating that elevated Cdc42 restricts lifespan via a mechanism independent of rDNA silencing.

**Fig. 2.**
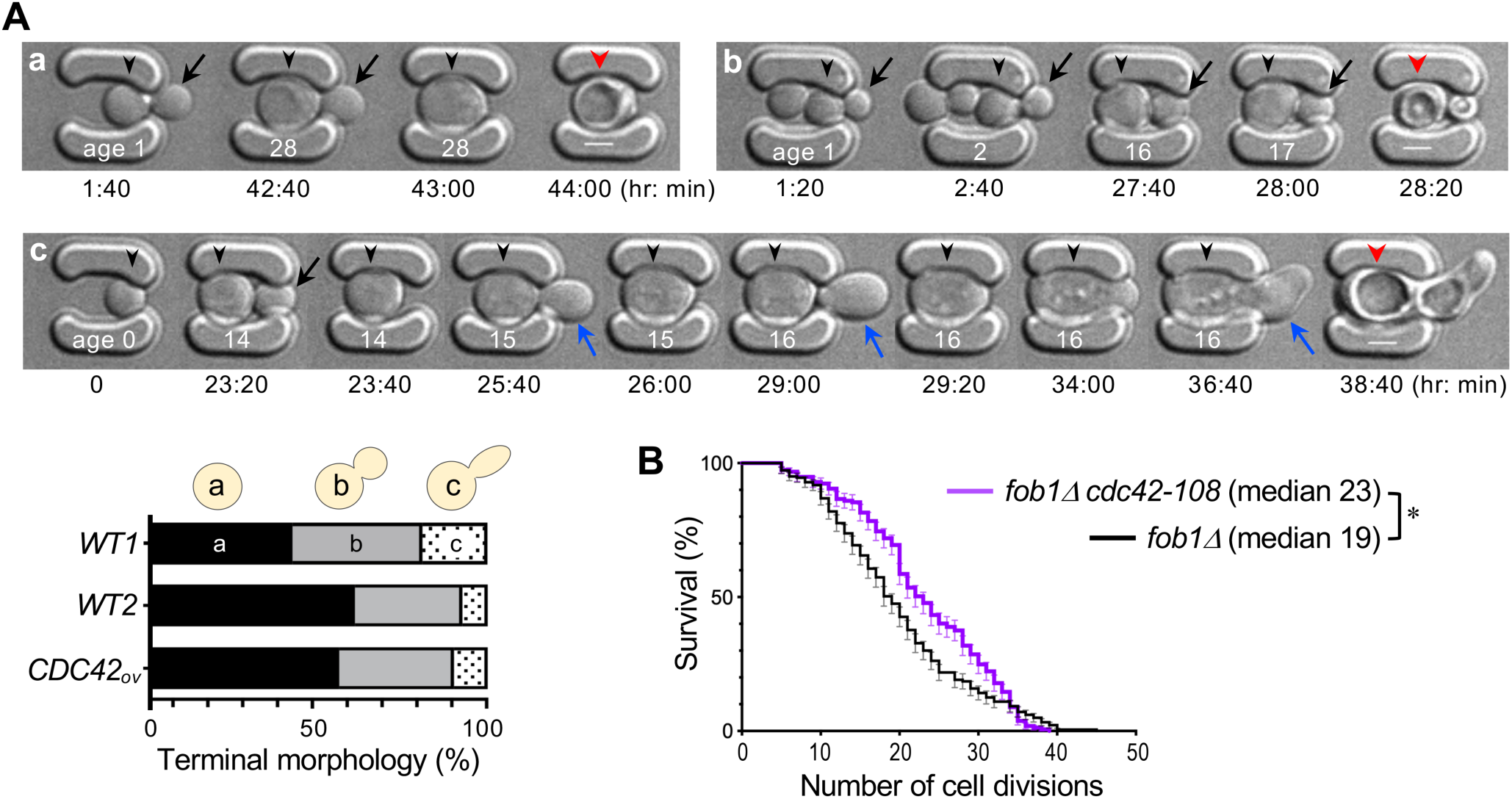
Cdc42 upregulation limits lifespan independently of rDNA desilencing **A.** Representative microfluidic images show different cell morphologies during aging: (a) a cell dying in the unbudded state, (b) a cell producing round buds (black arrows) until death, and (c) a cell forming elongated buds (blue arrows). Age and time points are shown in white and black letters. Scale bar: 3 μm. Quantification of distinct terminal cell shapes is shown below for WT1 (BY4741), and two isogenic strains, WT2 (HPY210) and *CDC42_OV_* (HPY3721) (n = 105∼110 per strain). **B.** Cell survival curves (mean ± SEM) are shown for *fob1Δ* (n = 183) and *fob1Δ cdc42-108* mutants (n = 157). Median lifespan is indicated; *, *p* < 0.05, Log-Rank test.

### Cdc42 is asymmetrically distributed during cell division through its association with endomembranes

We measured Cdc42-mCherry^SW^ fluorescence in mother and daughter pairs of WT across generations. These analyses showed that daughter cells generally inherit lower Cdc42 levels than their mothers, regardless of the mother’s age (Fig. 3A; see Cell 1, Fig. 1A). In rare cases (∼19%, n = 37), both mother and daughter cells died shortly after division. These daughter cells, which failed to have full lifespan, inherited high Cdc42 levels comparable to those of their mothers (Fig. 3B) and were similar in size to their mothers at division (see Cell 2 at t = 14:20, Fig. 1A). These observations suggest that Cdc42 accumulates in mother cells during asymmetric division and that lower Cdc42 levels in daughters are likely to help maintain their youthfulness.

**Fig. 3.**
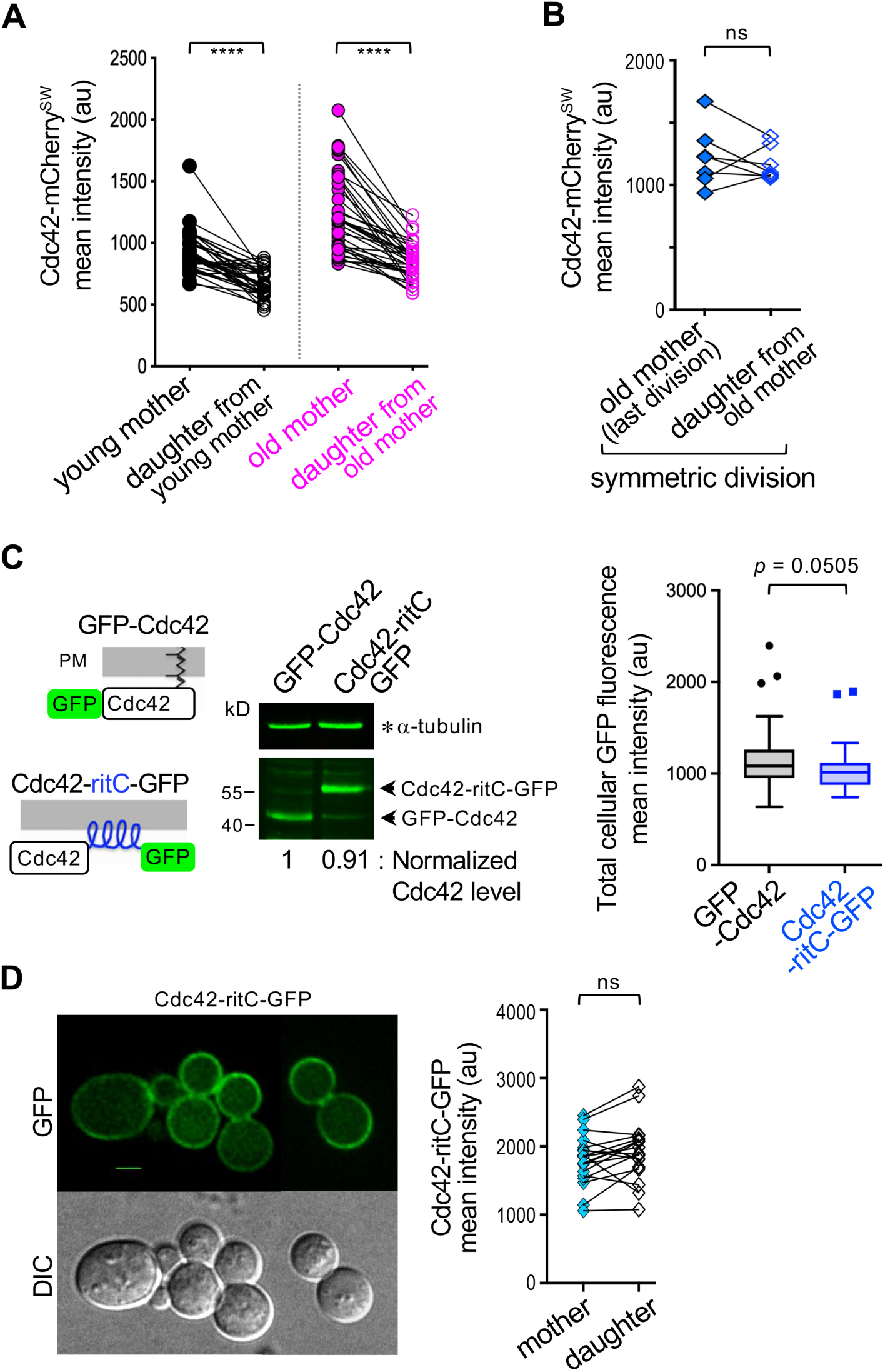
Asymmetric distribution of Cdc42 during cell divisions via endomembrane association **A.** Cdc42-mCherry^SW^ levels in each pair of young and old mothers and their daughters during M-cytokinesis (n = 37 pairs, each group). ****, *p* < 0.0001 from paired t-tests. Refer to Fig. 1B legend. **B.** Cdc42-mCherry^SW^ levels in mother and daughter pairs at the final division preceding death of both cells (see Cell 2 in Fig. 1A). n = 7 (out of 37 lineages); ns, *p* ≥ 0.05, paired t-test. **C.** Schematic diagrams of WT Cdc42 and Cdc42-ritC-GFP with the Rit amphipathic helix (in blue). Levels of GFP-Cdc42 and Cdc42-ritC-GFP were compared by immunoblotting extracts from cells (expressing each allele as the sole genomic copy) grown at 25°C to early log phase, using a monoclonal anti-GFP antibody and α-tubulin as a loading control. The mean GFP fluorescence in whole cells (mother and bud combined) of these strains, grown at 25°C, is shown in a Tukey plot (n = 72 per strain); ns, *p* ≥ 0.05, unpaired t-test. **D.** Localization of Cdc42-ritC-GFP at 25°C, and quantification in mother and bud compartments during M-cytokinesis. 20 representative pairs are shown. ns, *p* ≥ 0.05, paired t-test. Scale bar: 3 µm. See S2_Fig and S2_Movie.

To understand how Cdc42 is distributed asymmetrically, we considered two mechanisms: (1) retrograde transport of Cdc42 from the bud to the mother cell, as seen in protein aggregate segregation [23]; and (2) age-related changes in proteasome activity influencing protein turnover [39]. Retrograde transport occurs during apical growth [40], when Cdc42 is concentrated at the bud tip (see S1_Fig).

While aging might alter proteasome function and Cdc42 turnover, there is little evidence for different turnover rates between young mother cells and their daughters. Thus, neither mechanism fully explains Cdc42’s asymmetrical distribution.

We next examined whether Cdc42’s membrane association affects its distribution, since Cdc42 localizes to both the <u>plasma membrane (**PM**</u>) and endomembranes [27, 28, 30]. We confined Cdc42 to the PM using a *cdc42-ritC* mutant, in which its C-terminal CaaX motif is replaced with the amphipathic tail of mammalian Rit GTPase [41]. The *cdc42-ritC-GFP* strain grew normally at 24–27°C but poorly above 30°C, occasionally forming multiple buds simultaneously, as previously reported [41] (S2_Fig; S2_Movie). Cdc42-ritC-GFP levels were comparable to GFP-Cdc42; both were expressed from the endogenous *CDC42* promoter as the sole genomic copy, indicating unaffected protein stability (Fig. 3C). However, Cdc42-ritC-GFP lost its asymmetric localization, showing similar levels in mother and daughter cells at division (Fig. 3D). These results suggest that Cdc42’s association with and release from internal membranes are critical for its asymmetric distribution.

The *cdc42-ritC* mutant likely has a shortened lifespan. Because of abnormal cell shapes and a multi-budding phenotype of *cdc42-ritC-GFP* cells, we estimated lifespan by time-lapse imaging, instead of tracking successive mother cell divisions by microfluidic imaging or micromanipulation. Strikingly, most newborn daughter cells died after only a few divisions at 25°C (>90%, n = 100) (S2_Fig; S2_Movie), supporting the idea that Cdc42 asymmetry is critical for the full lifespan of daughter cells. However, in addition to defects in endomembrane association, *cdc42-ritC-GFP* cells frequently failed to establish cell polarity after a few divisions, as indicated by the appearance of large, round mother cells that stopped budding (S2_Fig; S2_Movie). These complex phenotypes of the *cdc42-ritC* mutant complicate directly linking its reduced lifespan to the protein’s symmetric distribution.

### Farnesylated Ydj1 interacts with Cdc42 and supports its asymmetry and stability

Because endomembrane association is necessary for Cdc42 partitioning, we investigated the possible role of Ydj1, a farnesylated Hsp40 tethered to the ER [42, 43]. Ydj1 is enriched in mother cells via an ER diffusion barrier and retains protein aggregates in aging cells [12]. As *ydj1Δ* mutants grow poorly at 30°C and are inviable at 37°C [42, 44] (see S3_Fig), we examined Cdc42-mCherry^SW^ localization in *ydj1Δ* cells at 27°C, a permissive temperature. In these cells, Cdc42 polarized normally during early cell cycle stages but became symmetrically distributed during division (Fig. 4A). We quantified Cdc42-mCherry^SW^ in the mother and bud compartments of large-budded *ydj1Δ* cells (Fig. 4B, left) and calculated asymmetry as the log of the mother/bud fluorescence ratio (Fig. 4B, right). These analyses show that *ydj1Δ* cells are defective in establishing Cdc42 asymmetry during division.

**Fig. 4.**
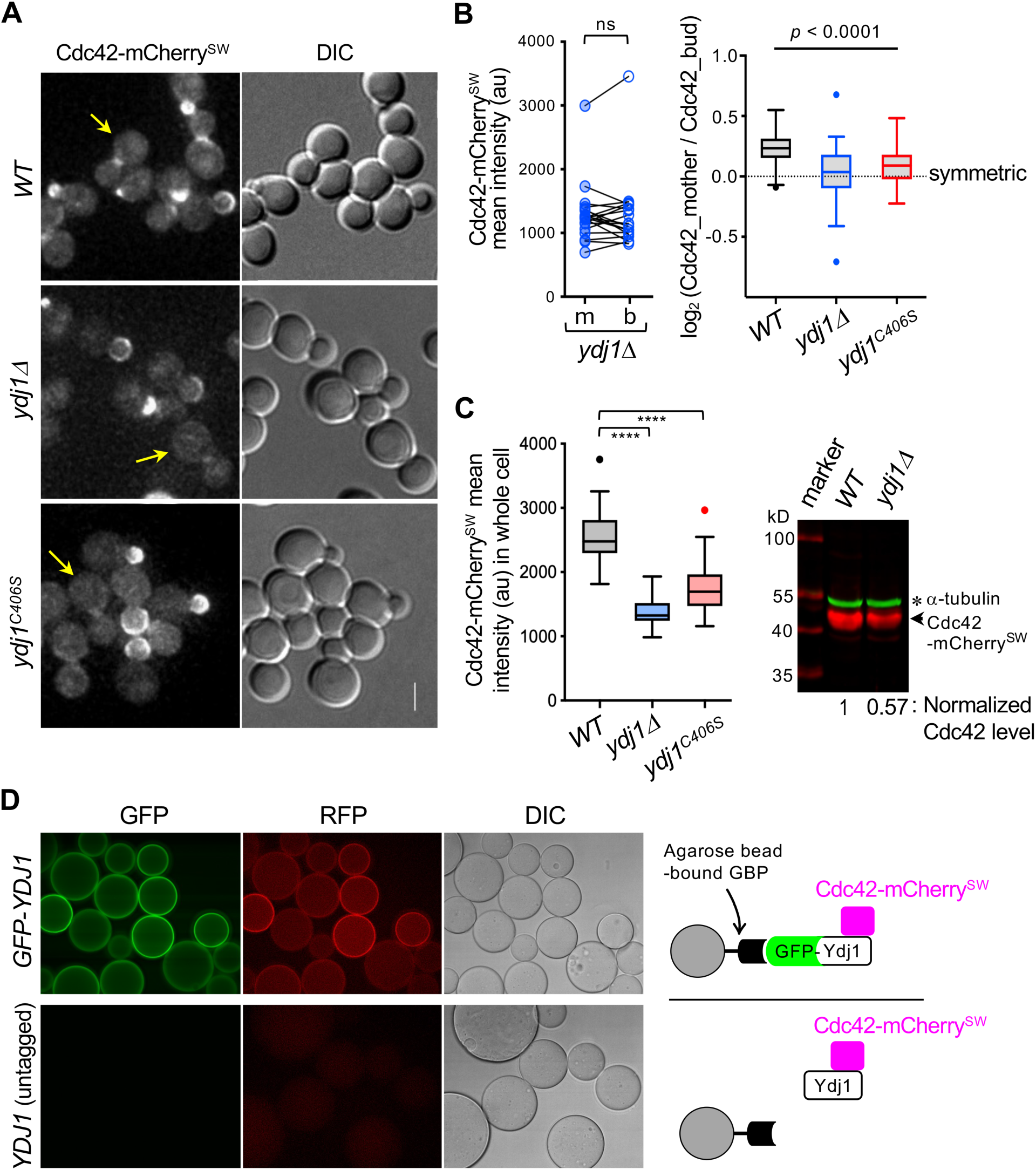
Farnesylated Ydj1 is required for maintaining Cdc42 levels and its asymmetric distribution. **A.** Localization of Cdc42-mCherry^SW^ in WT and *ydj1* mutants at 27°C. Arrows mark examples of large-budded cells used for Cdc42 quantification in **B** & **C**. Scale bar: 3 µm. See S3_Fig. **B.** Cdc42-mCherry^SW^ levels in mother (m) and bud (b) compartments of large-budded *ydj1Δ* cells, grown at 27°C. (left plot) Mean fluorescence intensities from 19 representative mother-bud pairs are plotted. ns, *p* ≥ 0.05, paired t-test. (right plot) The log_2_ mother-to-bud ratio of Cdc42-mCherry^SW^ mean intensity in WT and *ydj1* mutants (n = 52 per strain). The dotted line denotes a symmetric distribution of Cdc42 between mother and bud. ****, *p* < 0.0001 by one-way ANOVA. See also S3_Fig. **C.** Cdc42 levels in WT and *ydj1* mutants, grown at 27°C to mid-log phase. Mean fluorescence intensities of Cdc42-mCherry^SW^ in whole cells (mother and bud combined) are plotted. n = 57∼60 per strain; ****, *p* < 0.0001, unpaired t-tests. Immunoblotting shows Cdc42-mCherry^SW^ in each strain, detected using polyclonal anti-RFP antibodies, and α-tubulin, a loading control. See also S3_Fig. **D.** Association of Cdc42-mCherry^SW^ with GFP-Ydj1 detected by a visible IP assay (top panel). A control reaction used extracts containing untagged Ydj1 (bottom panel).

Since farnesylated Ydj1 diffuses more slowly in mother cells than in buds [12], we hypothesized that ER-anchored Ydj1 facilitates Cdc42’s asymmetric distribution during division. Supporting this, we found that Cdc42 asymmetry is partially reduced in the farnesylation-defective *ydj1(C406S)* mutant [45], though to a lesser extent than in *ydj1Δ* cells (Fig. 4B, right). Statistical analysis confirmed a significant association between farnesylated Ydj1 and Cdc42 asymmetry (one-way ANOVA, *p* < 0.0001). Additionally, Cdc42 protein levels were reduced in *ydj1Δ* and *ydj1(C406S)* mutants compared to WT, as shown by Cdc42-mCherry^SW^ fluorescence analysis in whole cells and by immunoblotting (about a two-fold reduction in *ydj1Δ* cells; Fig. 4C).

We next examined whether Cdc42 physically associates with Ydj1 using a <u>visible</u> <u>immunoprecipitation (VIP)</u> assay [46], which combines immunoprecipitation with fluorescence microscopy. In this assay, lysates from cells expressing GFP-Ydj1 and Cdc42-mCherry^SW^ were pulled down with <u>GFP-binding protein (GBP</u>)-conjugated beads [47]. Fluorescence microscopy showed that both GFP-Ydj1 and Cdc42-mCherry^SW^ were retained on the beads, while no signal was detected in control samples with untagged Ydj1 (Fig. 4D), demonstrating an in vivo interaction between Cdc42 and Ydj1.

Our study highlights Ydj1’s critical role in maintaining Cdc42 protein stability and its asymmetric distribution. Ydj1 functions along with Hsp70 and Hsp90 chaperones in protein folding and stabilization, including protein kinase maturation [48, 49]. Previous work showed that the farnesylation-defective *ydj1(C406S)* mutation affects the stability and maturation of the Hsp90 client Ste11, while mutations in the substrate-binding domain I (SBD I) of Ydj1 do not affect Ste11 accumulation [50]. To test if this also applies to Cdc42, we examined the *ydj1(L135S)* mutation in the SBD I peptide-binding pocket [51, 52]. A low-copy *ydj1(L135S)* plasmid rescued the temperature-sensitive growth defect of *ydj1Δ* mutants, as previously reported [51]. Cdc42 levels in *ydj1Δ* cells carrying either *L135S* or WT *YDJ1* plasmids were similar and higher than those in cells with *C406S* or an empty vector (S3_Fig, C). Quantification of Cdc42 levels separately in mother and bud compartments showed that the *L135S* mutation did not significantly alter Cdc42’s asymmetric distribution (WT vs. *L135S*, *p* > 0.05), unlike the *C406S* mutation (WT vs. *C406S*, *p* < 0.001) (S3_Fig). Together with analysis of *ydj1(C406S)* cells (see Fig. 4, B & C), these results suggest that farnesylation of Ydj1 is critical for both Cdc42’s stability and asymmetric distribution, likely via ER tethering.

In contrast to our results at 27°C, previous studies reported higher Cdc42 levels in *ydj1Δ* cells compared to WT at 30°C and after shifting to 37°C [53, 54]. The reason for these different findings is unclear, but it may be due to temperature-dependent stress responses. At 30°C or above, *ydj1Δ* mutants exhibit impaired growth and increased expression of Hsp70 and Hsp90 [42, 44, 49], which may facilitate Cdc42 stabilization and proper folding at higher temperatures. Thus, the increased Cdc42 observed in *ydj1Δ* mutants under these conditions might reflect indirect compensatory mechanisms rather than the direct effect of losing Ydj1. Further studies are required to clarify these mechanisms under different conditions.

### Summary and Limitations

This study demonstrates that Cdc42 levels increase as yeast cells age and that reducing these levels can extend lifespan. Cdc42 is distributed unevenly between mother and daughter cells during division.

Daughter cells inheriting high Cdc42 levels, similar to their mothers, often die within a few divisions, suggesting that proper asymmetric distribution is crucial for their full replicative lifespan. However, whether a specific Cdc42 threshold determines proliferative capacity remains unclear. Our mutational and biochemical analyses show that Cdc42’s asymmetric distribution depends on its association with endomembranes and interaction with the chaperone Ydj1. Notably, Ydj1 is necessary for both maintaining proper Cdc42 levels and facilitating its asymmetric distribution. We propose that Ydj1 anchors Cdc42 to the ER in mother cells, leading to its uneven segregation and the establishment of age-related asymmetry. However, many questions remain, including how farnesylated Ydj1 anchors Cdc42 in mother cells and whether Ydj1 collaborates with Hsp90 to help Cdc42 maturation remains unknown. This study also did not directly address how Ydj1 impacts this process during aging. Further studies, such as manipulating Cdc42 levels directly or testing additional *CDC42* and *YDJ1* alleles with specific defects, could help elucidate these mechanisms.

While our data suggest that Cdc42 promotes senescence independently of rDNA silencing loss, how Cdc42 upregulation drives aging remains unclear. Cdc42 is not one of the “long-lived asymmetrically retained proteins” typically associated with accumulated cellular damage [55].

Interestingly, many mother cell-enriched proteins whose depletion extends lifespan are normally turned over, indicating that aging or senescence is not solely due to damage accumulation [56, 57]. We speculate that repeated asymmetric divisions lead to Cdc42 accumulation in mother cells, eventually impairing cellular function through aberrant signaling and thus limiting proliferative capacity. While our study raises new questions, it reveals a chaperone-dependent mechanism for Cdc42 partitioning during asymmetric division, which likely contributes to cellular aging. This may be a conserved mechanism in other asymmetrically dividing cells.

## MATERIALS AND METHODS

### Yeast Strains, Growth Conditions, and Plasmids

Standard yeast genetics methods, DNA manipulation, and growth conditions were used [58]. Yeast strains were cultured in rich YPD medium (yeast extract, peptone, dextrose) or in the appropriate synthetic medium containing 2% dextrose as a carbon source. All fusion proteins were expressed from their native promoters on the chromosomes, except for GFP-Ydj1, which was expressed from a plasmid for visible immunoprecipitation assays. Unless otherwise indicated, yeast cultures for imaging and protein preparation were grown at 27°C. The low-copy plasmids pRS315-YDJ1, pRS315-ydj1(C406S), pRS315-ydj1(L135S) (CEN, *LEU2*), and pRS416-GFP-YDJ1 (CEN, *URA3*) were generous gifts from D. M. Cyr (University of North Carolina) and W. Schmidt (University of Georgia) [45, 52]. Yeast strains used are listed in S1_Table.

### Microscopy and Microfluidic Imaging

Cells were grown in an appropriate synthetic medium overnight and then freshly subcultured for 3–4 hrs in the same medium. Live-cell imaging, including microfluidics-based imaging, was performed using an inverted microscope (Ti-E; Nikon) fitted with a 100x/1.45 NA Plan-Apochromat Lambda oil immersion objective lens, a 60x/1.4 NA Plan Apochromat Lambda oil immersion objective lens, and DIC optics (Nikon), FITC/GFP, and mCherry/Texas Red filters from Chroma Technology, an Andor iXon Ultra 888 electron-multiplying charge-coupled device (EM CCD) (Andor Technology), Sola Light Engine (Lumencor) solid-state illumination, and the software NIS Elements (Nikon).

Static fluorescence images (Figs. 3C & 3D, 4A–4C, and S3_Fig) were captured using a 100x/1.45 NA objective lens (11 z stacks, 0.4 μm step) with cells either mounted on a 2% agarose slab or a glass-bottomed dish (MatTek) containing an appropriate synthetic medium, as previously described [59, 60]. Due to aberrant cell shapes and a multi-budding phenotype, the lifespan of *cdc42-ritC-GFP* cells was estimated using long-term time-lapse imaging. Freshly subcultured cells were seeded at a low density in a glass-bottomed dish (MatTek), and images were acquired with a 60x/1.4 NA objective lens (9 z-stacks, 0.5 μm steps) using DIC optics every 25 min for approximately 16 hrs. Representative timepoints are shown in S2_Fig and S2_Movie. The slab or dish was put directly in a stage top chamber (Okolab) set to 27°C or at room temperature (24–25°C), as marked in figure legends or figures.

Microfluidics setup and growth conditions are essentially the same as previously described [32], except imaging temperature was maintained at 27°C. Microfluidic devices were fabricated using polydimethylsiloxane (PDMS) by adopting a design [32] that allows simultaneous imaging of multiple strains through independent passages. Microfluidics-assisted time-lapse imaging was performed using an inverted widefield fluorescence microscope (Ti-E; Nikon) equipped with a 60x/1.4 NA objective lens (5 z stacks, 0.5 μm step) and DIC optics (see above). In general, bright-field images were recorded every 20 min throughout the entire experiment at multiple XY positions, and fluorescence images were captured for the initial 3∼4 hrs and a few hours after 12 hrs, 26 hrs, and 50 hrs, except as indicated. In some cases (Fig. 1A), fluorescence images were captured every 20 min with a minimum exposure throughout the entire experiment.

### Image Processing and Analysis

Images were processed by importing nd2 files using the NIH ImageJ [61] with Bio-Format importer plugin. Mean fluorescence intensities of Cdc42-mCherry^SW^ were quantified by defining a region of interest (ROI) in single-focused z-stack images using the ImageJ oval selection tool only in cells during the late M phase or cytokinesis; i.e., when no polarized Cdc42 signals were detectable at the bud periphery or the incipient bud site (e.g., cells marked with white arrowheads in Fig. 1A). Since asymmetric distribution of Cdc42 between mother and bud compartments was established at this cell cycle stage (see S1_Fig), Cdc42 levels at this cell cycle stage were quantified throughout this study. These values were compared between young and old ages (of the same cell lineages) or between a mother and her daughter by averaging the mean intensity of Cdc42-mCherry^SW^ in the first three divisions (young mothers) or last three divisions (old mothers) (Figs. 1A–1B & 3A). Cdc42 levels were similarly quantified in mother and bud compartments separately in large-budded cells from the exponential growth culture at the indicated temperature (Figs. 3D & 4B; S3_Fig). Cdc42 asymmetry was plotted as the log_2_ of the Cdc42-mCherry^SW^ fluorescence ratio in each compartment (mother/bud) (Fig. 4B; S3_Fig). Cdc42-mCherry^SW^ levels in whole cells (mother and bud combined) of WT and mutants were compared by quantifying mean fluorescence intensity using single best-focused z slices, except by defining an ROI covering both a mother and its bud using the ImageJ freehand tool (Fig. 4C; S3_Fig).

Cellular levels of Cdc42-ritC-GFP and GFP-Cdc42 were quantified similarly in whole cells from images of cells in the exponential growth culture at 27°C (Fig. 3C).

To make figures, fluorescence images were deconvolved by the Iterative Constrained Richardson-Lucy algorithm (NIS Elements) and cropped at selected time points, preferentially at cell division or soon after division. The same setting of brightness/contrast was applied to adjust images in multiple panels within the same figure.

### Lifespan Estimation

Lifespan was estimated by counting the number of cell divisions observed in microfluidic images starting from initial loading until cell death or senescence. Cell death was identified by abrupt cell shrinkage or lysis seen in DIC images, a sudden and complete loss of fluorescence signal, or the appearance of strong autofluorescence throughout the cell. Cells were considered at senescence or near senescence if their cell cycle length increased sharply (by more than ∼6 hrs) without subsequent division for 8–12 hrs. Lifespan estimation from microfluidic imaging sometimes underrepresents extremely long-lived cells compared to micromanipulator-based assays, mainly due to rare cell loss or crowding events at later timepoints. Unlike in micromanipulator-based assays, not all initially loaded cells were newborn daughter cells, so the total cell divisions observed were less than the replicative lifespan. Nonetheless, relative differences in the median number of cell divisions between strains were reproducible, especially since multiple strains were imaged simultaneously using a chamber with multiple flow passages. Cells that died after fewer than five divisions were excluded from lifespan analysis, as certain daughter cells born from very old mothers did not reach their full lifespan potential [62].

### TCA Protein Extraction and Immunoblotting

WT and mutant strains were cultured in YPD medium at the temperatures specified in the figure legends and harvested at OD_600_ 0.6–0.8. Whole cell extracts were prepared by precipitation with 10% trichloroacetic acid (TCA), as previously described [63]. Protein precipitates were then resuspended in 100 mM Tris (pH 11.0) and 3% SDS, heated at 90°C for 5 min, and briefly centrifuged to remove cell debris. Proteins were separated by SDS-PAGE on 12.5% polyacrylamide gels, and western blots were visualized using the LI-COR Odyssey system (LI-COR Biosciences, Lincoln, Nebraska) with the following antibodies: mouse monoclonal anti-Cdc42 antibody (clone 28-10) (EMD Millipore, MABN2485), rabbit polyclonal anti-RFP antibodies (Rockland, 600-401-379), mouse monoclonal anti-GFP antibodies (GFP-ID2-s) (DSHB, University of Iowa), and mouse monoclonal anti-alpha tubulin antibody (clone 12G10) (DSHB, University of Iowa). Secondary antibodies used were Alexa Fluor^®^ 680 goat anti-rabbit IgG (ThermoFisher Scientific, A32734) or IRDye^®^ 800CW goat anti-mouse IgG (LI-COR Biosciences, 926-32210). Total protein levels in each sample were normalized using Ponceau S-stained membranes. To ensure that α-tubulin, used as a loading control, was present at similar levels relative to total protein in both WT and mutant strains, non-specific cross-reactive bands were analyzed for comparison.

### Visible Immunoprecipitation (VIP) Assay

VIP assays were performed using a total of 80 OD_600_ units of HPY4079 (*CDC42-mCherry^SW^ ydj1Δ*) cells transformed with either pRS416-GFP-YDJ1 or pRS315-YDJ1 (control), as previously described [60] with modifications. Crude cell lysates were prepared in PBS (PBS, pH 7.4, 0.5 mM EDTA) containing a protease inhibitor cocktail (Research Products International) and 0.1 mM PMSF, followed by centrifugation at 14,000 rpm for 15 min at 4°C to isolate the pellet fraction (P10). Proteins were solubilized from P10 using 300μL of extraction buffer (PBS, pH 7.4, 200 mM NaCl, 0.5 mM EDTA, 2% Triton) at 4°C for 1 h and then subjected to pull-down assays with GFP-Trap beads (gta-10, Chromotek). After washing, beads were mounted on an agarose slab and imaged on a Nikon TiE inverted microscope with a 40x/0.75 Plan Fluor objective at room temperature. Static images were captured using DIC, FITC/GFP, or mCherry/Texas Red filters (5 z-stacks, 0.5μm step), and single z-slices were selected for figure preparation.

### Statistical Analysis

Data analysis was performed using Prism 10 (GraphPad Software). To determine statistical differences between two sets of cell survival data, the Gehan-Breslow-Wilcoxon test and Log-rank (Mantel-Cox) test were used. For comparison of Cdc42 levels between young and old cells in the same lineage or between mothers and their daughters, paired t-tests were performed to determine statistical significance: ns (not significant) for *p* ≥ 0.05; \**p* < 0.05, \*\**p* < 0.01, \*\*\**p* < 0.001, and \*\*\*\**p* < 0.0001. Statistical significance of the Cdc42-mCherry^SW^ mean intensity differences between mother-bud pairs among WT and mutants was determined by one-way ANOVA analysis. For comparison of Cdc42-mCherry^SW^ in whole cells (mother and bud combined) between WT and mutants, unpaired t-tests were performed. For box graphs, the Tukey method was used to create whiskers (for minimum and maximum data points), a box with a line (for the upper, lower quartiles, and median), and to mark outliers.

## ACKNOWLEDGMENTS

We are grateful to M. Peter and the Department of Biology at ETH Zurich for their generous support with microfluidics and for hosting a short-term visit to the Institute of Biochemistry, ETH Zurich. We also thank K. Kozminski, D. Drubin, W. Schmidt, D. G. Cyr, S. Martin, and D. Lew for yeast strains and plasmids. The authors declare no competing financial interests.

## AUTHOR CONTRIBUTIONS

PJK: Conceptualization, Investigation, Methodology, Resources, Supervision, Visualization, Writing – original draft, Writing – review & editing HM: Investigation, Visualization, Writing – review & editing SSL: Methodology, Resources, Writing – review & editing HOP: Conceptualization, Methodology, Visualization, Funding acquisition, Project administration, Supervision, Writing – original draft, Writing – review & editing

## Abbreviations

ERCs: extrachromosomal rDNA circles
PM: plasma membrane
ER: endoplasmic reticulum
WT: wild type

## Supporting Information

**S1_Fig. Quantification of Cdc42-mCherry^SW^ levels in growing buds**

Mean fluorescence intensities of Cdc42-mCherry^SW^ within growing buds are compared. A split septin ring (labeled with Cdc3-GFP) indicates the onset of cytokinesis. Arrows mark the same bud developing and producing a daughter; an arrowhead marks a daughter cell from the previous division. Scale bar: 3 µm. n = 32∼40 per group; ****, *p* < 0.0001, Welch’s t-test.

S2_Fig. Growth phenotype and time-lapse imaging of *cdc42-ritC* mutant.

**A.** Growth phenotypes of *cdc42-ritC, cdc42-ritC-GFP,* and isogenic wild-type (WT) strains on YPD plates at 27°C, 30°C, and 34°C.

**B.** Time-lapse images of *cdc42-ritC-GFP* cells at 25°C. Selected time points are shown every 50 min for about 16 hours. Colored asterisks mark daughter cells (d1–d3) originating from the same mother cell (black arrows). Red arrows denote cell death. Note: Daughter cell d2 continued to grow large without typical cell death signs. See S2_Movie.

S3_Fig. Additional characterization of *ydj1* mutants

A. Growth phenotypes of *ydj1Δ*, *ydj1(C406S)*, and isogenic WT strains on YPD plates at 27°C, 30°C, and 37°C.

B. Growth phenotypes of *ydj1Δ* cells carrying either WT YDJ1, ydj1(C406S), ydj1(L135S), or pRS315 plasmid on SC-Leu plates at 27°C, 30°C, and 37°C.

C. Mean fluorescence intensities of Cdc42-mCherry^SW^ in whole cells (mother and bud combined) of the *ydj1Δ CDC42-mCherry^SW^*strain carrying each plasmid, grown at 27°C (n = 78∼86 per strain). ****, *p* < 0.0001; *, *p* = 0.0333, by unpaired Welch’s t-test.

D. The log_2_ mother-to-bud ratio of Cdc42-mCherry^SW^ mean intensity (mean ± SEM) in *ydj1Δ* cells carrying each plasmid, grown at 27°C (n = 22–40 per group). *p* = 0.0005 by ordinary one-way ANOVA; ns, *p* ≥ 0.05; **, *p* < 0.01; and ***, *p* < 0.001 by Welch’s t-test.

**S1_Movie. Microfluidic imaging of WT cells expressing Cdc42-mCherry^SW^**

Images were captured every 20 min at 27°C using an inverted widefield microscope (TiE, Nikon) with a 60x/ 1.4 NA objective lens, and fluorescence images were deconvolved. The movie shows frames for 56 hr 20 min of Cell 1, with selected time points shown in Fig. 1A. The display rate is 8 frames per sec (fps).

**S2_Movie. Time-lapse imaging of Cdc42-ritC-GFP cells**

Images were captured every 25 min at 25°C using an inverted widefield microscope (TiE, Nikon) with a 60x/ 1.4 NA objective lens and DIC optics. The movie shows frames every 50 min for 15 hr 50 min. The display rate is 8 frames per sec (fps). An example of cropped images is shown in S2_Fig.

**S1_Table. Yeast strains used in this study**

