## Supplementary figures and images for "Cdc42 Partitioning by Chaperone Ydj1 During Asymmetric Division and Aging in Yeast"

### Supplemental Data 1

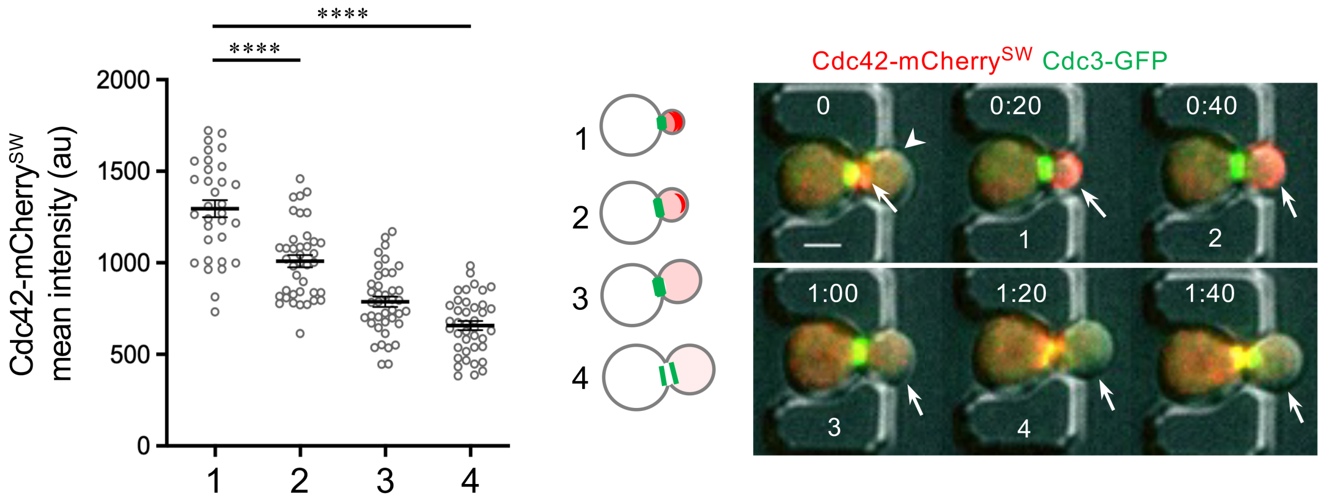

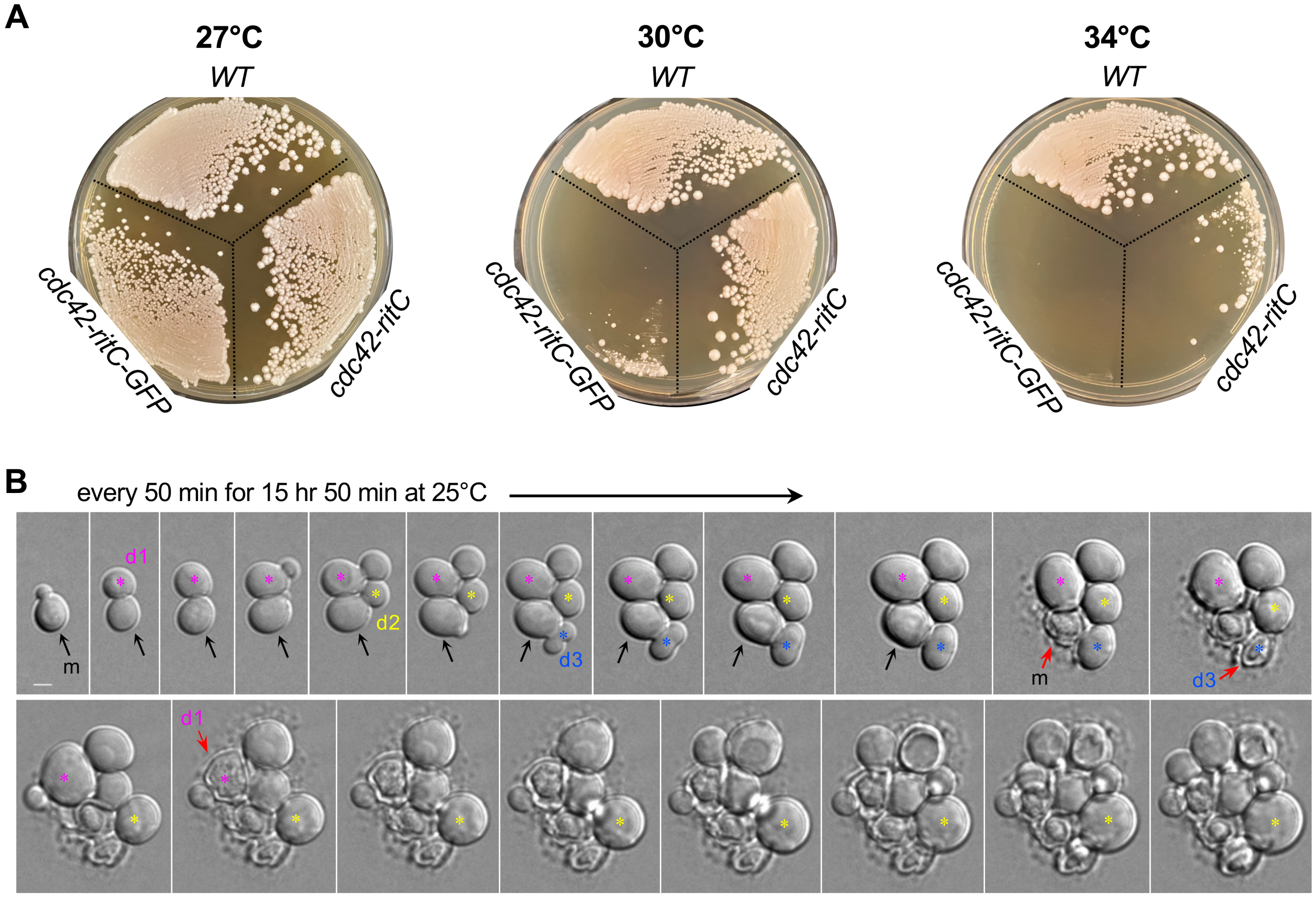

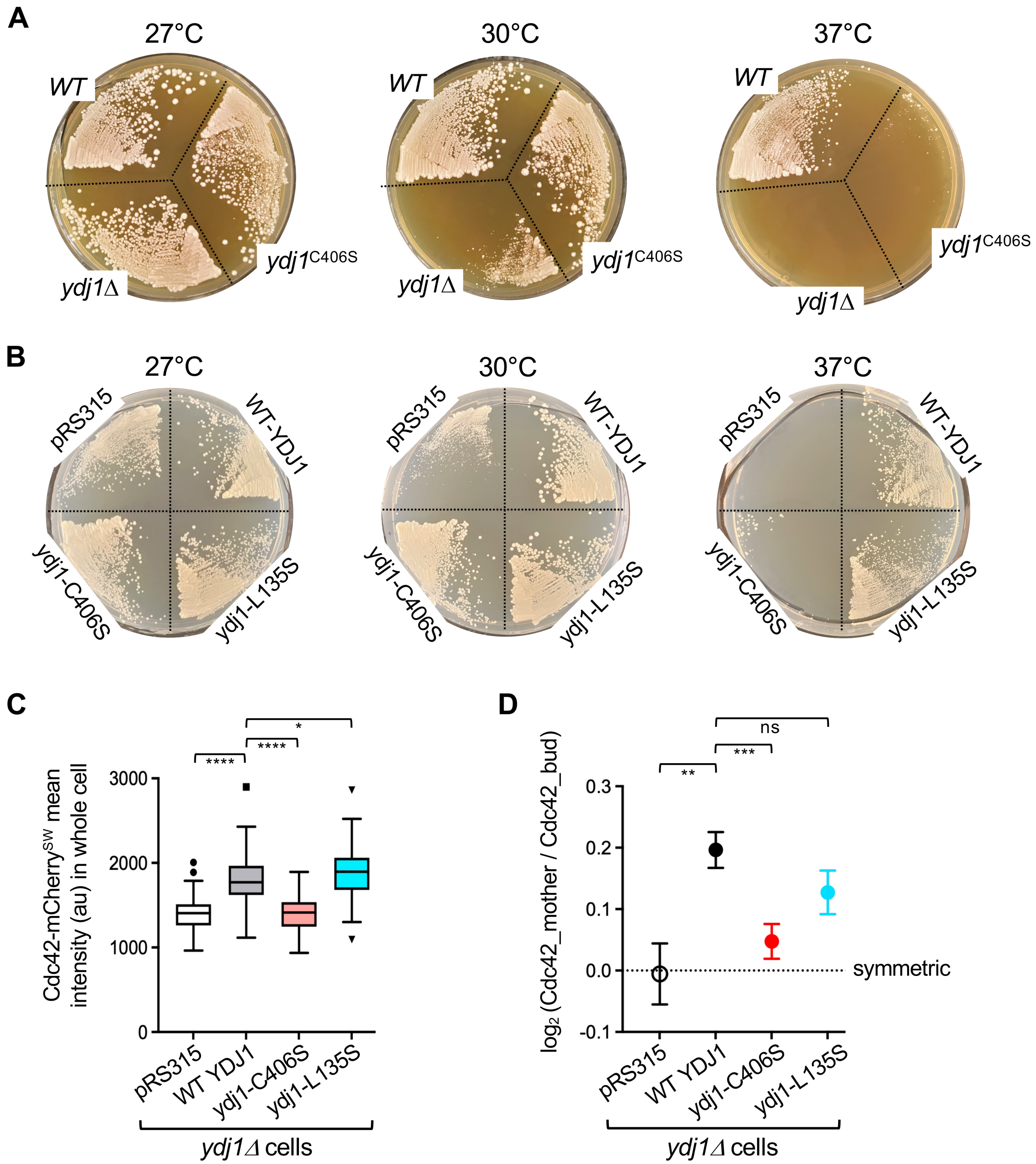
