## Supplementary material for "Cdc42 Partitioning by Chaperone Ydj1 During Asymmetric Division and Aging in Yeast": S1_Table

**Table S1. Yeast strains used in this study**

| Strain[a] | Relevant Genotype | Source/Comments |
|---|---|---|
| BY4741[@] | *MAT***a** *his3Δ1 leu2Δ0 met15Δ0 ura3Δ0* | Open Biosystems |
| YSC13[@] | $P_{CDC42}$*-GFP-cdc42-ritC-kanMX* | [1] |
| YSC11[@] | *MAT***a** *cdc42-ritC-GFP-kanMX* | [1] |
| DDY1300[@] | *MAT***a** *his3Δ200 leu2-3,112 lys2-801am ura3-52 CDC42:LEU2* | [2] |
| DDY1312[@] | *MAT***a** *cdc42-108*$^{R147A,\ E148A,\ K150A}$*:LEU2* | [2] |
| DDY4890[@] | *MAT***a** *CDC42-mCherry*$^{SW}$*:HIS3* | [3] |
| YWS2110[@] | *MAT***a** *his3Δ1 leu2Δ0 met15Δ0 ura3Δ0 ydj1*$^{C406S}$ | [4] |
| HPY3740[@] | *MAT***a** *CDC42-mCherry*$^{SW}$*::HIS3 CDC3-GFP:LEU2* | This study |
| HPY3881[@] | *MAT***α** *fob1Δ::KAN* | Open Biosystems |
| HPY3963[@] | *MAT***α** *cdc42-108:LEU2 fob1Δ::KAN met15 ura3 leu2* | Segregant of DDY1312 X HPY3881 |
| HPY4079[@] | *MAT***a** *CDC42-mCherry*$^{SW}$*:HIS3 ydj1Δ::KAN* | This study |
| HPY4092[@] | *MAT***α** *CDC42-mCherry*$^{SW}$*:HIS3 ydj1*$^{C406S}$ | Derived from YWS2110 |
| HPY4148[@] | *MAT***a** *cdc42-108*$^{R147A,\ E148A,\ K150A}$*:LEU2 cdc42-108*$^{R147A,\ E148A,\ K150A}$*:URA3* | This study[b] |
| HPY210[#] | *MAT***a** *his3-Δ200 leu2-Δ1 lys2-801 trp1-Δ63 ura3-52* | [5] |
| HPY3721[#] | *MAT***a** *cdc42::TRP1 GFP-CDC42(8X)-URA3 CDC3-mCherry:LEU2* | [6] |
| HPY3215[#] | *MAT***a** *cdc42::TRP1* $P_{CDC42}$*-GFP-CDC42:URA3* | Segregant of DLY13920 [7] |

[a] Strains marked with [@] are derived from DDY1300 or BY4741 (both in S288C background); and strains marked with [#] are isogenic with HPY210, derived from YEF473 [8].

[b] Copy of *cdc42-108*$^{R147A,\ E148A,\ K150A}$ was obtained from DDY1312 by genomic PCR and cloned on pRS306. *Eco*RV-digested DNA was used to integrate an additional copy of *cdc42-108*$^{R147A,\ E148A,\ K150A}$ into the *URA3* locus of DDY1312.
